# Estimating the genetic parameters of yield-related traits under different nitrogen conditions in maize

**DOI:** 10.1101/2022.08.05.502993

**Authors:** Semra Palali Delen, Gen Xu, Jenifer Velazquez-Perfecto, Jinliang Yang

## Abstract

Understanding the genetic basis responding to nitrogen (N) fertilization in crop production is a long-standing research topic in plant breeding and genetics. Albeit years of continuous efforts, the genetic architecture parameters, such as heritability, polygenicity, and mode of selection, underlying the N responses in maize remain largely unclear. In this study, about *n* = 230 maize inbred lines were phenotyped under high N (HN) and low N (LN) conditions for two consecutive years to obtain six yield-related traits. Heritability analyses suggested that traits highly responsive to N treatments were less heritable. Using publicly available SNP genotypes, the genome-wide association study (GWAS) was conducted to identify *n* = 231 and *n* = 139 trait-associated loci (TALs) under HN and LN conditions, respectively, and *n* = 162 TALs for N-responsive (NR) traits. Furthermore, genome-wide complex trait Bayesian (GCTB) analysis, a method complementary to GWAS, was performed to estimate genetic parameters, including genetic polygenicity and the mode of selection (*S*). GCTB results suggested that the NR value of a yield component trait was highly polygenic and that four NR traits exhibited negative correlations between SNP effects and their minor allele frequencies (or the *S* value < 0) — a pattern consistent with negative selection to purge deleterious alleles. This study reveals the complex genetic architecture underlying N responses for yield-related traits and provides insights into the future direction for N resilient maize development.

## Introduction

Nitrogen (N), as a fundamental macronutrient, is a major constituent of proteins, nucleic acid, and metabolites and is critical for the high yielding of crops (1). Since the 1960s, subsequent to the Green Revolution, due to the Haber-Bosch process, inorganic N fertilizers became increasingly available for crop production, especially in maize, where about 20% of the N fertilizers was applied for maize production (2; 3). However, inefficient N usage causes ammonia emission to the environment, accounting for a considerable proportion of fine particulate matter pollution (i.e., PM2.5) and reducing human population life span (4). Meanwhile, N runoff imposes substantial adverse effects on natural ecosystems, such as reduced water quality and impaired soil health. Therefore, understanding the plant response to N in crop production is crucial for human health, food security, and environmental sustainability and is a long-standing research topic in plant breeding and genetics.

To identify N-responsive genetic loci, many QTL studies were performed, resulting in a number of trait-associated QTLs under different N conditions (5; 6) or QTLs for different N-related traits, i.e., grain N yield, N remobilization, and post-silking N uptake (7; 8). Recently, as the technical advances, genetic studies for N-related traits shifted from QTL mapping to GWAS (9; 10), leading to high-resolution mapping results. For example, a recent GWAS using 411 maize inbred lines under optimum and low N conditions detected about 80 significant SNPs and 136 putative candidate genes (11). These N-related QTLs and trait-associated SNPs provide opportunities to investigate the fate of the deleterious alleles — the alleles can potentially affect fitness under different N conditions. During the recent maize improvement process, an excess of the mutational load was enriched in even elite maize inbred lines (12). However, it is largely unclear how many alleles contribute to NR traits and what is the mode of selection on these alleles, including potentially deleterious alleles, in affecting N responses.

In the current study, by employing two complementary approaches — GWAS and GCTB (Genome-wide Complex Trait Bayesian analysis), we analyzed yield-related traits collected under low N (LN) and high N (HN) conditions (i.e., trait *per se*) as well as the transformed N-responsive (NR) traits. We found higher heritability for most traits *per se* under HN than LN and identified 1,292 trait-associated SNPs in total that locate in 481 genomic regions. Inferring from genome-wide non-zero effects SNPs, including not only significant GWAS SNPs but also SNPs with minor effects, GCTB results suggested the yield-related NR traits were highly polygenic and that NR traits were more likely under negative selection (13). The complex genetic architecture revealed from this study, especially for the NR traits, provides guidelines for further genome-enabled selection modeling and N resilient maize development.

## Materials and Methods

### Plant materials and field experimental design

In this experiment, a subset (*n* = 226 genotypes) of the maize diversity panel (14) was planted in a rain-fed experimental field followed commerical maize. For the N treated plots, urea (dry fertilizer) as a source of N was applied at the rate of 120 lbs/acre before planting. The field experiment was conducted using an incompletely randomized block design in two consecutive field seasons (2018-2019). For each replication of a treatment, the field was split into four blocks by plant height and maturity (i.e., tall/early, tall/late, short/early, short/late). Each block was further subdivided into three sub-blocks. Within each sub-blocks, two hybrid varieties B73×Mo17 and B37×Mo17 were planted randomly as check plants (see also (15; 16)).

### Phenotypic data collection

From each two-row plot, three mature ears were harvested from the representative plants. These harvested ears were dried in the oven at 37°*C* for three days to decrease the moisture content. Harvested ears were hand-shelled to prevent kernel loss. After shelling, the kernels and cobs were kept separately with proper barcoded labels. From the cobs, cob diameter (CD), cob length (CL), and cob weight (CW) were manually measured. The total kernel weight (TKW) of each ear was measured from the collected kernels. And then, 20 representative kernels were selected to measure 20 kernel weight (20KW). Finally, the kernel count (KC) was computed using TKW divided by average kernel weight.

### Best linear unbiased prediction (BLUP) and N-responsive trait calculation

To obtain the best linear unbiased prediction (BLUP) values of each genotype, we fitted a linear mixed model by treating the genotype, year, replication, block, sub-block, and genotype by year interaction as random effects. For each N treatment, the BLUP values were calculated separately using an R package “lme4” (17).

In the model,

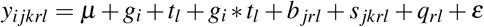

where *y*_*i jkrl*_ is the phenotypic value of the *i*^*th*^ genotype evaluated in the *k*^*th*^ sub-block of the *j*^*th*^ block of *r*^*th*^ replicate nested within the *l*^*th*^ year; *μ* is the overall mean; *g*_*i*_ is the random effect of the *i*^*th*^ genotype; *t*_*l*_ is the random effect of the *l*^*th*^ year; *g*_*i*_ * *t*_*l*_ is the random effect of the *i*^*th*^ genotype with the *l*^*th*^ year interaction; *b*_*jrl*_ is the random effect of the *j*^*th*^ block of the *r*^*th*^ replicate within the *l*^*th*^ year; *s* _*jkrl*_ is the random effect of the *k*^*th*^ sub-block of the *j*^*th*^ block of the *r*^*th*^ replicate within the *l*^*th*^ year; *q*_*rl*_ is the random effect of the *r*^*th*^ replicate nested within the *l*^*th*^ year; *ε* is the random residual error.

The N-responsive (NR) traits were calculated from the BLUP values using the equation (18):

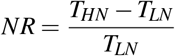

where *T*_*HN*_ and *T*_*LN*_ are the BLUP values for a given trait measured from HN and LN field conditions.

### Broad sense heritability calculation

The broad-sense heritability (*H*^2^) of yield-related traits was calculated using the equation as:

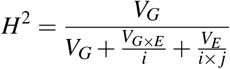

where *V*_*G*_ is the genotypic variance; *V*_*E*_ is the environmental variance of different years; and *V*_*G*×*E*_ is the variance of genotype by year interaction; *i* = 2 is the number of years and *j* = 2 is the number of replications per year.

### Genome-wide Association Study (GWAS)

The SNP genotype of the maize diversity panel was downloaded from maize HapMap3 (19) with AGPv4 coordinates. After filtering out SNPs with minor allele frequency (MAF) < 0.05 and missing rate < 0.3 among the 226 lines phenotyped in this study, approximately 21 million SNPs were retained.

In GWAS, we employed the QK model that considers both population structure (Q) and kinship relatedness (K) to control for multiple levels of confounding effects (20; 21). In the model,

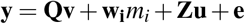

where **y** is a vector of BLUP value for a given trait (or the NR trait); **Q** is the design matrix of the population structure (i.e., the principle components); **v** is the vector of the fixed subpopulation effect; **w**_**i**_ is a vector of the *i*^*th*^ SNP genotype; *m*_*i*_ is the fixed SNP effect to be estimated by an iterative procedure; **Z** is the covariance matrix or the kinship matrix of inbred lines; **u** is the vector of breeding values to be predicted (random effect); **e** is the vector of the random residual error.

In the analysis, the **Q** matrix was the first three principal components calculated from genome-wide SNPs using PLINK 1.9 software (22). And the **Z** matrix was computed using GEMMA (v 0.98.3) software with option “4” (23). The above model was then implemented to estimate significant SNP effects for each trait using GEMMA (23). The threshold for the significant association SNPs was set to 1.2 × 10^−6^ (1/n, *n* = 769,690 is the number of independent SNPs with MAF ≥ 5%) according to the method developed previously (15). From the GWAS results, significant genomic loci were determined by considering a 100 kb window upstream and downstream of the significant SNPs. Overlapping regions were merged, and these regions were defined as trait-associated loci (TAL).

### Genome-wide Complex Trait Bayesian (GCTB) analysis

Genome-wide Complex Trait Bayesian (GCTB-BayesS) approach, which is based on Bayesian multiple regression mixed linear models (24), was performed to estimate genetic architecture parameters of yield-related traits, including polygenicity (number of non-zero SNPs) and selection coefficient (the joint distribution between the variance of SNP effects and minor allele frequencies). Default options were selected with the following MCMC settings: “–chain-length = 1,010,000, –burn-in = 10,000”. In the analysis, *n* = 834,975 independent SNPs (MAF ≥ 1%) were used (25), which was determined by using PLINK 1.9 (22) with the “indep-pairwise” option (window size 100 kb, step size 100, *r*^2^ ≥ 0.1).

## Results

### Phenotypic evaluation of diverse maize lines under different N conditions

A subset of the maize diversity panel (*n* = 226 lines) was planted in a replicated field trial under high N (HN) and low N (LN) conditions according to an incomplete block design in 2018 and 2019 (see **Materials and Methods**). From the harvested mature ears, six yield-related traits were manually measured, including three cob-related traits (cob diameter, CD; cob length, CL; and cob weight, CW) and three kernel-related traits (20 kernel weight, 20KW; kernel count, KC; and total kernel weight, TKW). For these traits, the best linear unbiased prediction (BLUP) values were calculated separately for each N condition. Besides traits *per se*, we also calculated N-responsive (NR) traits from the BLUP values following a previous method (26) (see **Materials and Methods** and **Table S1**).

As expected, most of these yield-related traits exhibited significantly larger BLUP values in HN than LN conditions, except for CD (**Figure 1A**). TKW, a trait most closely related to yield (12), showed the most striking differences from 39.9 g in LN to 46.9 in HN (Paired t-test, *P*-value = 1.9 × 10^−69^) and 97.8% of the lines exhibited positive N responses (**Figure 1B**). Similarly, the BLUP values for the other two kernel-related traits were also significantly improved from LN to HN, i.e., from 4.6 g to 4.8 g for 20KW (Paired t-test, *P*-value = 4 × 10^−4^) and from 174 to 196 for KC (Paired t-test, *P*-value = 3.9 × 10^−32^). And 63.3% and 88.6% of the inbred lines positively responded to the elevated N levels for 20KW and KC, respectively (**Figure 1B**). For cob-related traits, the phenotypic differences between HN and LN were relatively minor; for example, there were no significant differences for CD between LN and HN (Paired t-test, *P*-value = 0.75).

**Figure 1.**
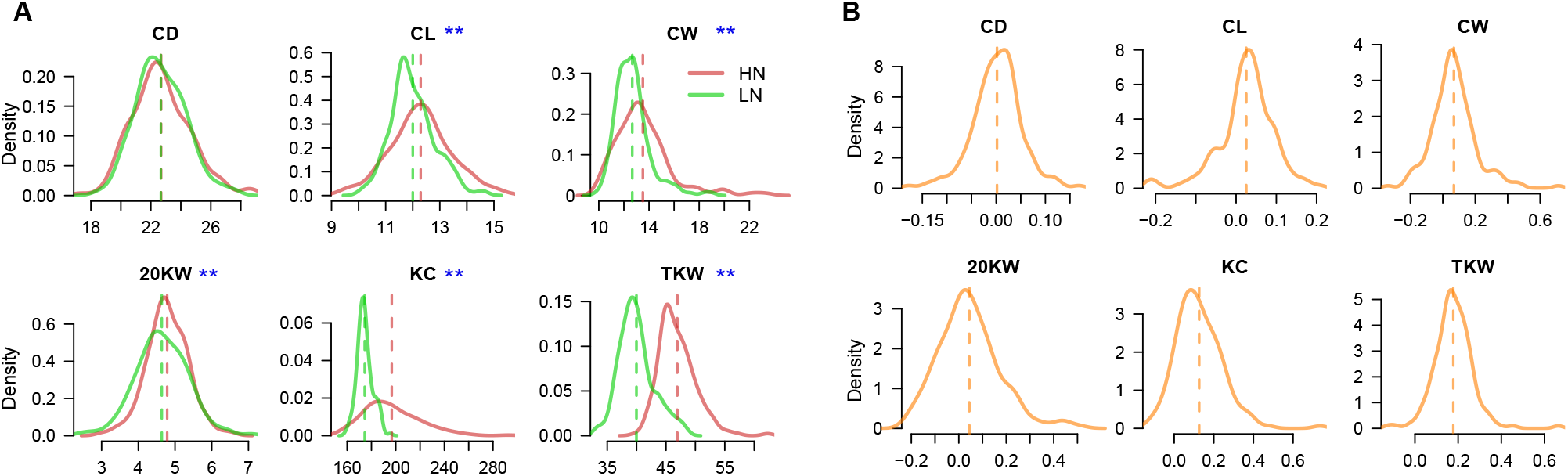
Cob- and kernel-related traits under different N conditions. (**A**) Density plots of the phenotypic traits in low N (LN) and high N (HN) fields. The red and green vertical dashed lines indicate the mean values of each trait. The blue asterisks indicate the traits show significant differences between HN and LN conditions (Paired t-test, *P*-value < 0.01). (**B**) The distributions of N-responsive (NR) traits. Orange dashed lines denote the mean values.

### Strong N-responsive traits are less heritable

The broad-sense heritability (*H*^2^) of these yield-related traits was estimated separately for each N condition (see **Materials and Methods**). Generally, we found these traits showed higher levels of heritability in HN than LN fields (**Figure 2A**), except for TKW (*H*^2^ = 0.38 in HN and *H*^2^ = 0.42 in LN), suggesting the environmental effects were less dominant or the data were more repeatable under HN conditions. For cob-related traits, CD (*H*^2^ = 0.81 in HN and *H*^2^ = 0.74 in LN) was more heritable than CL (*H*^2^ = 0.69 in HN and *H*^2^ = 0.53 in LN) and CW (*H*^2^ = 0.54 in HN and *H*^2^ = 0.37 in LN); while for kernel-related traits, 20KW (*H*^2^ = 0.73 in HN and *H*^2^ = 0.53 in LN) exhibited the highest heritability compared to KC (*H*^2^ = 0.47 in HN and *H*^2^ = 0.33 in LN) and TKW. Additionally, we found the cob-related traits, on average, were more heritable than kernel-related traits, regardless of the N conditions (**Figure 2A**). Interestingly, the levels of heritability negatively correlated with proportions of inbreds with NR values > 0 (**Figure 1B**), or ratios of inbreds positively responding to N treatments, under both N conditions (**Figure 2B**). These results are consistent with the view that more fitness-related traits, i.e., traits strongly responsive to N treatments, are less heritable (27).

**Figure 2.**
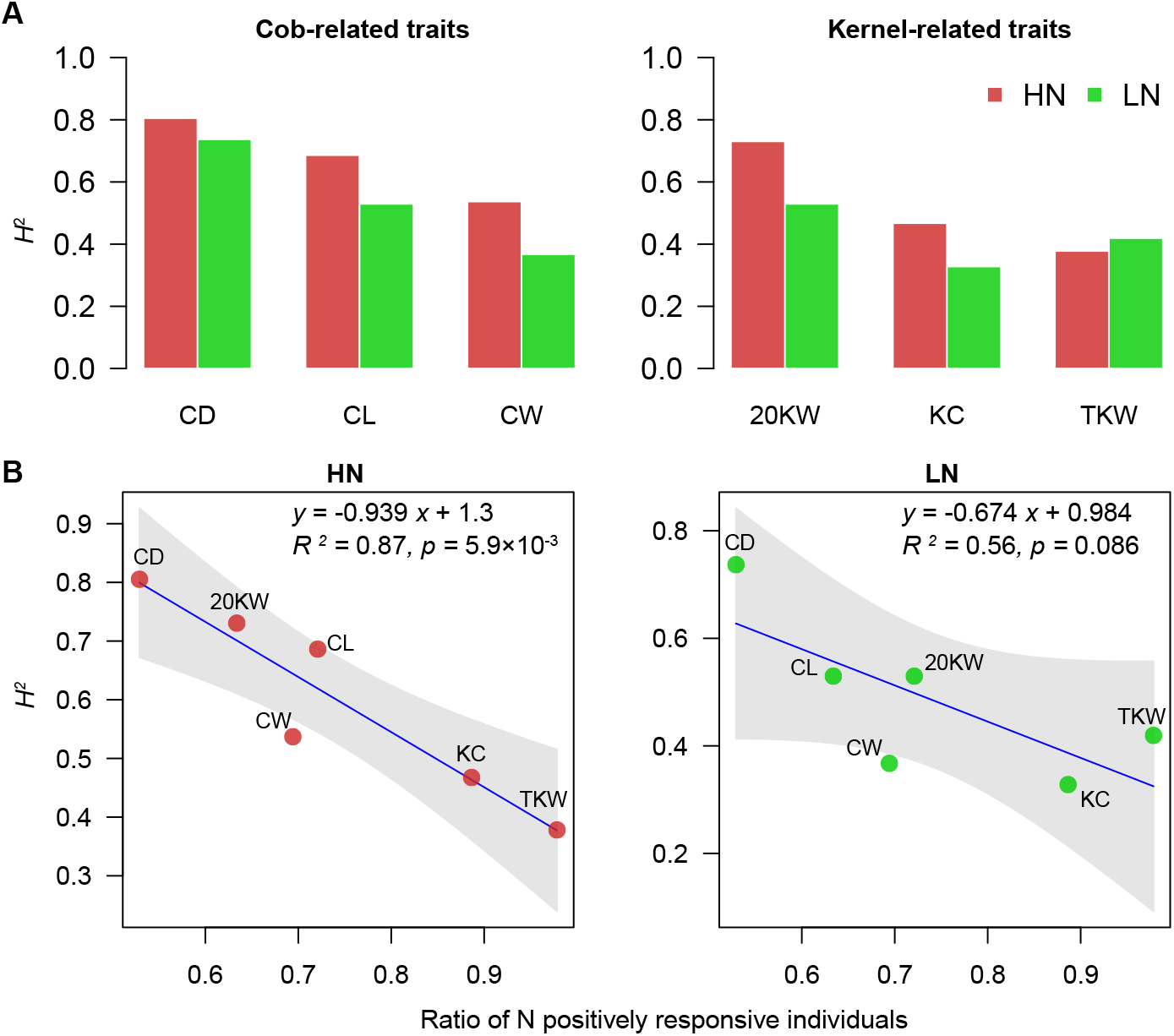
Heritability estimation and correlation analysis with N-response. (**A**) Heritabilities of yield-related traits under different N conditions. (**B**) Correlation analysis between heritability and N-responsive value. Solid blue line indicates the linear regression and the grey shaded area denotes the 95% confidence interval.

### Comparing GWAS signals under different N conditions

We then conducted GWAS for the six yield-related traits *per se* as well as the transformed NR traits by fitting a linear mixed model using 21 million SNPs (see **Materials and Methods**). In the GWAS model, the first three principal components were fitted as the fixed effects and the genetic relatedness computed from genome-wide SNPs as the random effects. To control for the false discovery rate (FDR), the modified Bonferroni-adjusted threshold was determined as 1.2 × 10^−6^ based on *n* = 769, 690 independent SNPs (28; 29). As a result (see **Figure S1, S2** for the quantile-quantile (Q-Q) plots), a total of 1,292 SNPs hitting 481 unique genomic regions were identified as the trait-associated loci (TALs, see **Table S2 - S3** for GWAS results).

We compared the shared TALs by different traits and treatments. For HN traits, in total, we identified 231 TALs (**Figure 3A**), *n* = 25 of which were detected for at least two yield-related traits (**Figure 3B**) — more than expected by chance (permutation test, *P*-value = 0.001). Such a large number of overlapping signals were not found for LN (*n* = 3 shared TALs, **Figure S4**) and NR (*n* = 6 shared TALs, **Figure S5**) traits (**Figure 3C-D**). Note that many shared TALs were between KC and TKW, likely because KC was not a directly measured trait but calculated from TKW and 20KW. Comparatively, very few overlapped TALs were identified for the same trait under different N conditions (**Figure S3**).

**Figure 3.**
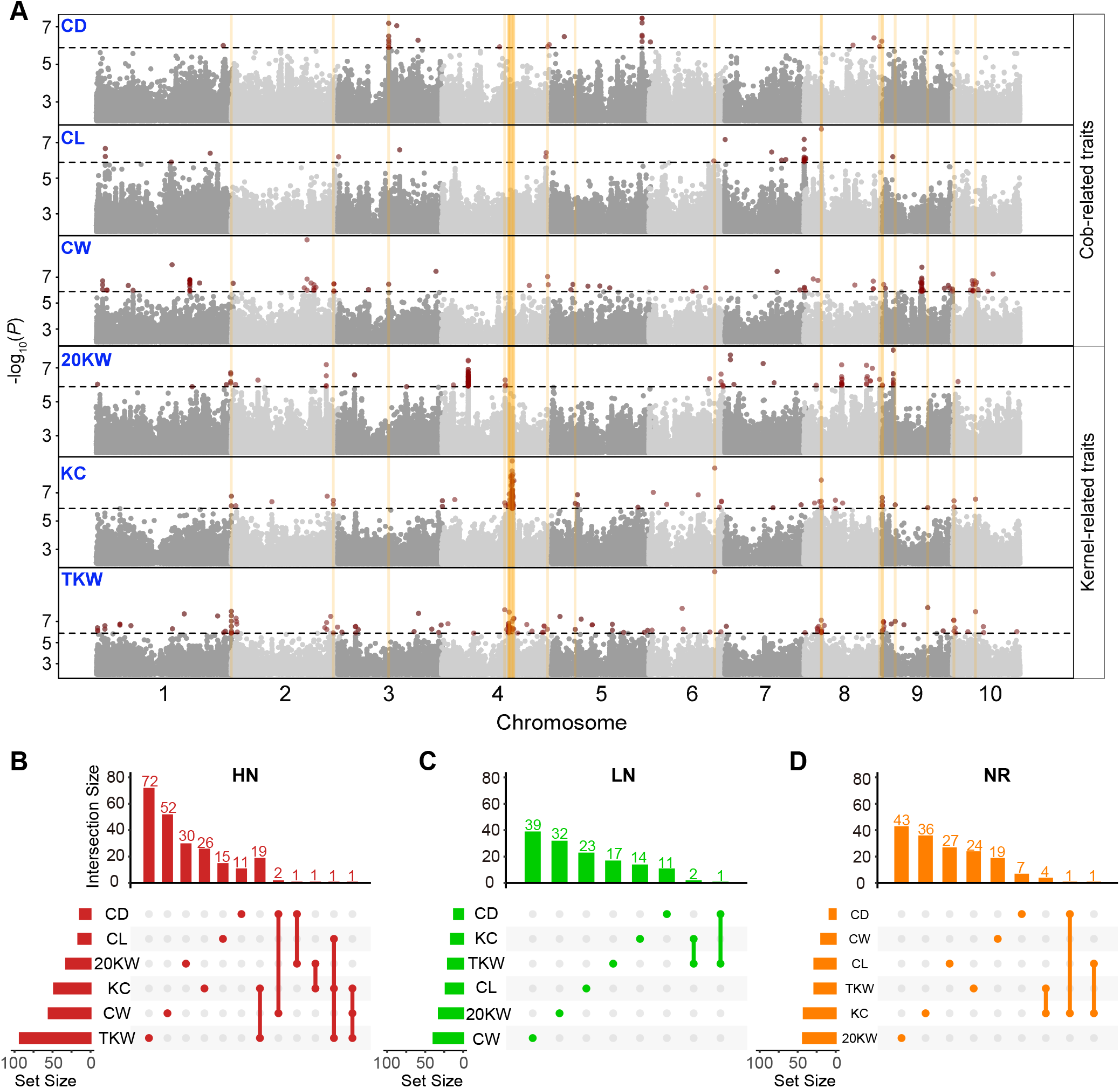
GWAS results for cob- and kernel-related traits under different nitrogen conditions. (**A**) Stacking Manhattan plot of cob- and kernel-related traits under HN conditions. The black horizontal dashed line indicates the GWAS threshold. Each red dot above the threshold represents the SNP significantly associated with a trait. The vertical orange lines indicate the overlapped trait-associated loci (TALs). (**B-D**) Overlapping results of TALs for HN (**B**), LN (**C**), and NR traits (**D**). Numbers on top of the barplots indicate the number of unique (only dots) and shared (dots and lines) TALs.

### Estimating genetic architecture parameters for traits *per se* and N-responsive traits

In addition to GWAS, we fitted the *per se* and NR traits to a Bayesian-based model implemented in GCTB (24). This method allows the simultaneous estimation of the genetic architecture parameters, such as polygenicity (*π*, the percentage of non-zero effect SNPs), variance of BLUP values due to SNPs (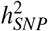, SNP-based heritability), and the mode of selection (*S*, a proxy using the relationship between variance of SNP effect and MAF). Using relatively independent SNPs with MAF > 1%, we estimated genetic parameters to compare the genetic architecture for the cob and kernel-related traits as well as for *per se* and NR traits (see **Materials and Methods**).

For trait *per se* under HN and LN conditions, we observed no significant differences in the polygenicity (**Figure 4A**), with an average of *n* = 906 SNPs (or *π* =0.1%) exhibiting non-zero effects; but the polygenicity among traits showed a large variation, ranging from *n* = 66 SNPs for CW in HN to *n* = 1,885 SNPs for CD in LN. In particular, for the NR trait of TKW, the number of non-zero effect SNPs elevated to *n* = 40,145 (or *π* =4.8%), suggesting TKW — a key yield component trait — was under complex genetic control in responding to changed N conditions. The 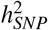 values for HN and LN traits were largely in line with the broad sense heritabilities estimated previously with the field data (**Figure 4B**). However, some abnormal values were detected, such as a small 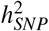 value for CL trait in LN, which was likely due to dominance or epistasis mode of inheritance playing an important role as our model considered only additive effect. Or simply because of imperfect model convergence. We also estimated the 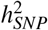 for NR traits and found, in general, kernel-related NR traits (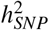 ranging from 0.55 ± 0.15 to 0.73 ± 0.1) were more heritable than cob-related NR traits (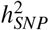 ranging from 0.3 ± 0.23 to 0. 73 ± 0.13). Finally, these estimated SNP effects and their allele frequencies in the population allowed us to infer the mode of selection. As pointed out by Zeng et. al., (24), *S* = 0 indicates selective neutral, while *S* > 0 and *S* < 0 suggest the positive and negative selection. Our results revealed that S values of four NR traits (CD, CL, CW, and 20KW), two HN traits (CW and TKW), and one LN trait (CW) were significantly smaller than zero (**Figure 4C**), indicating negative selection may be taken into effect to maintain the large effect deleterious SNPs in low frequencies, especially in responding to changed N conditions.

**Figure 4.**
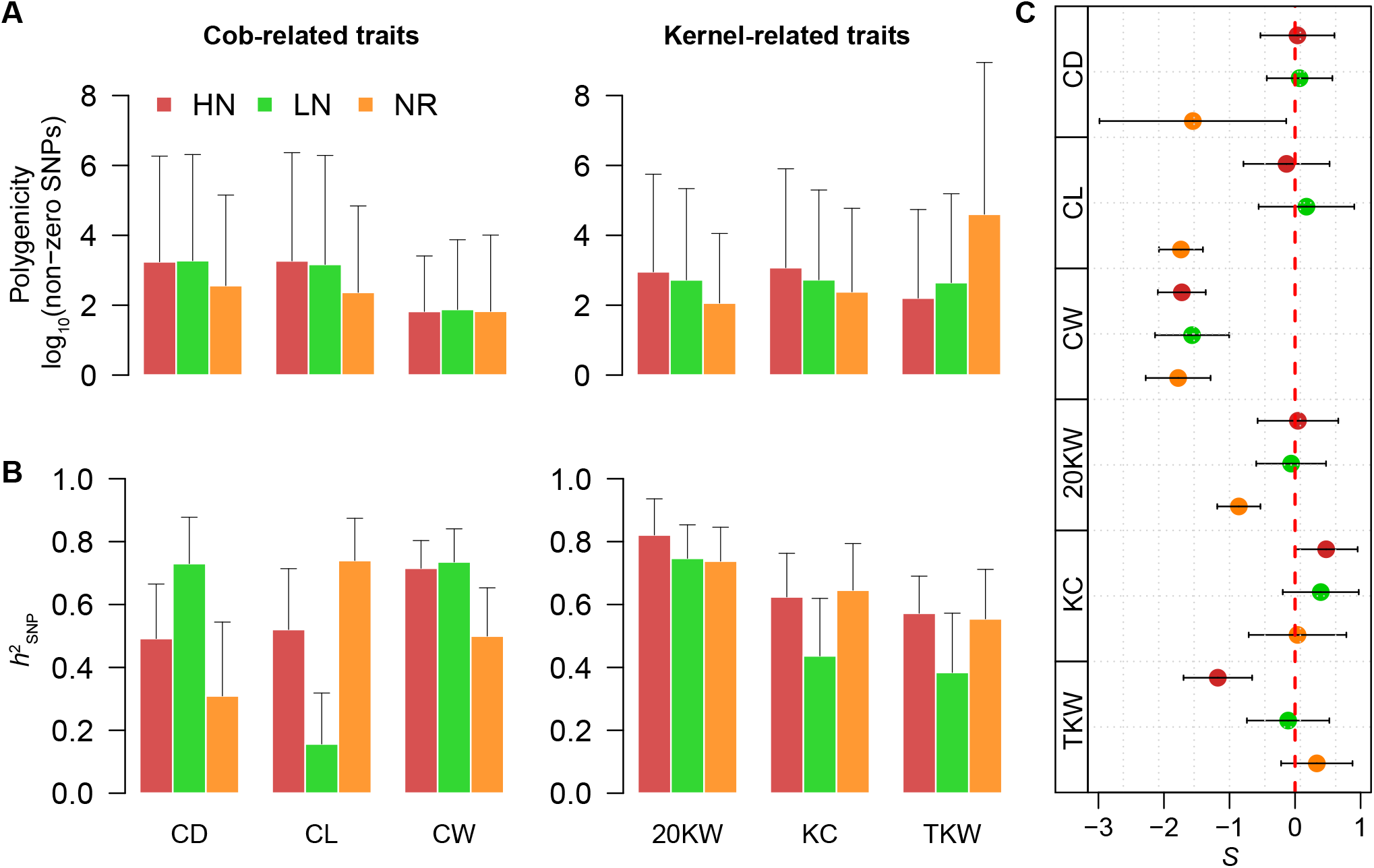
Genetic architecture parameters estimated for HN, LN and NR traits. (**A**) Genetic polygenicity measured by number of non-zero effect SNPs. (**B**) Phenotypic variance explained by SNPs under a additive model. (**C**) The relationships between SNP effects and MAFs of the non-zero effect SNPs.

## Discussion

In this study, we characterized six yield-related traits under high and low N conditions for two consecutive years and analyzed the data with two complementary approaches — GWAS (to estimate significant effect SNPs) and GCTB (to detect non-zero effect SNPs and infer other genetic parameters). We identified 1,292 GWAS signals located within *n* = 481 genomic regions or TALs. Many of these TALs were repeatedly detected for different traits under the same N conditions, but very few TALs were shared for the same trait under different N conditions, likely because genotype by N interaction plays an important role in controlling phenotypic variation. In addition to the GWAS SNPs, non-zero effect SNPs estimated from GCTB provided a proxy for evaluating genetic polygenicity. Results suggested that most of these yield-related traits are highly polygenic. In particular, we found the N-responsive trait of TKW was controlled by the highest number of non-zero SNPs (i.e., more than *n* = 4 × 10^4^ SNPs across the genome), consistent with the view that genetic basis for N responses in crop yield is highly complex (30). Heritability estimation from the field data suggested that traits highly responsive to N treatment tend to be less heritable, further confirming the genetic complexity of N responses for yield-related traits.

In the GCTB result, we detected most of the NR traits exhibiting negative S values, suggesting large effect SNPs for NR traits tend to be rare in the population. It is likely because these rare SNPs were deleterious and, therefore, were maintained in low frequencies to increase the plant fitness in responding to changed N conditions. N, as one of the significant macronutrients for crop development, its composition in the soil varies spatially and temporally (31; 32). Therefore, it is not surprising to expect plant breeding over the past 60 years since the Green Revolution or natural selection on a longer time scale has affected the patterns of deleterious alleles in responding to the N availability. However, the limitations of the current statistical methods (i.e., high false discovery rates and imperfect model convergence) prevent us from pinning down the individual deleterious locus accurately. Note that these genetic parameters were rough estimations derived from the posterior distributions. The point estimations were associated with large standard errors. To get more accurate results, a larger population or more sophisticated statistical approaches are warranted.

## Data and code availability

The code used for the analyses can be accessed through the GitHub repository (https://github.com/jyanglab/Genetics-parameters-for-N-related-traits).

## Acknowledgements

This work is supported by the Agriculture and Food Research Initiative Grant number 2019-67013-29167 from the USDA National Institute of Food, the National Science Foundation under award number OIA-1557417 for Center for Root and Rhizobiome Innovation (CRRI), and Agriculture and the University of Nebraska-Lincoln Start-up fund. This work was conducted using the Holland Computing Center of the University of Nebraska-Lincoln, which receives supports from the Nebraska Research Initiative.

## Author contributions

J.Y. designed this work. S.P.D. and J.V.-P. generated the data. S.P.D., G.X., and J.Y. analyzed the data. S.P.D., G.X., and J.Y. wrote the manuscript.

## Supplemental Material

### Supporting Tables

**Table S1. The best linear unbiased prediction (BLUP) values of yield-related traits**. (https://github.com/jyanglab/Genetics-parameters-for-N-related-traits/blob/main/supp%20tables/Stable1_phenotype.xlsx)

**Table S2. GWAS SNPs for cob- and kernel-related traits under different N conditions**. (https://github.com/jyanglab/Genetics-parameters-for-N-related-traits/blob/main/supp%20tables/Stable2_Significant_SNPs_from_GWAS.xlsx)

**Table S3. Trait-associated loci for cob- and kernel-related traits under different N conditions**. (https://github.com/jyanglab/Genetics-parameters-for-N-related-traits/blob/main/supp%20tables/Stable3_GWAS_Loci.xlsx)

### Supporting Figures

**Figure S1.**
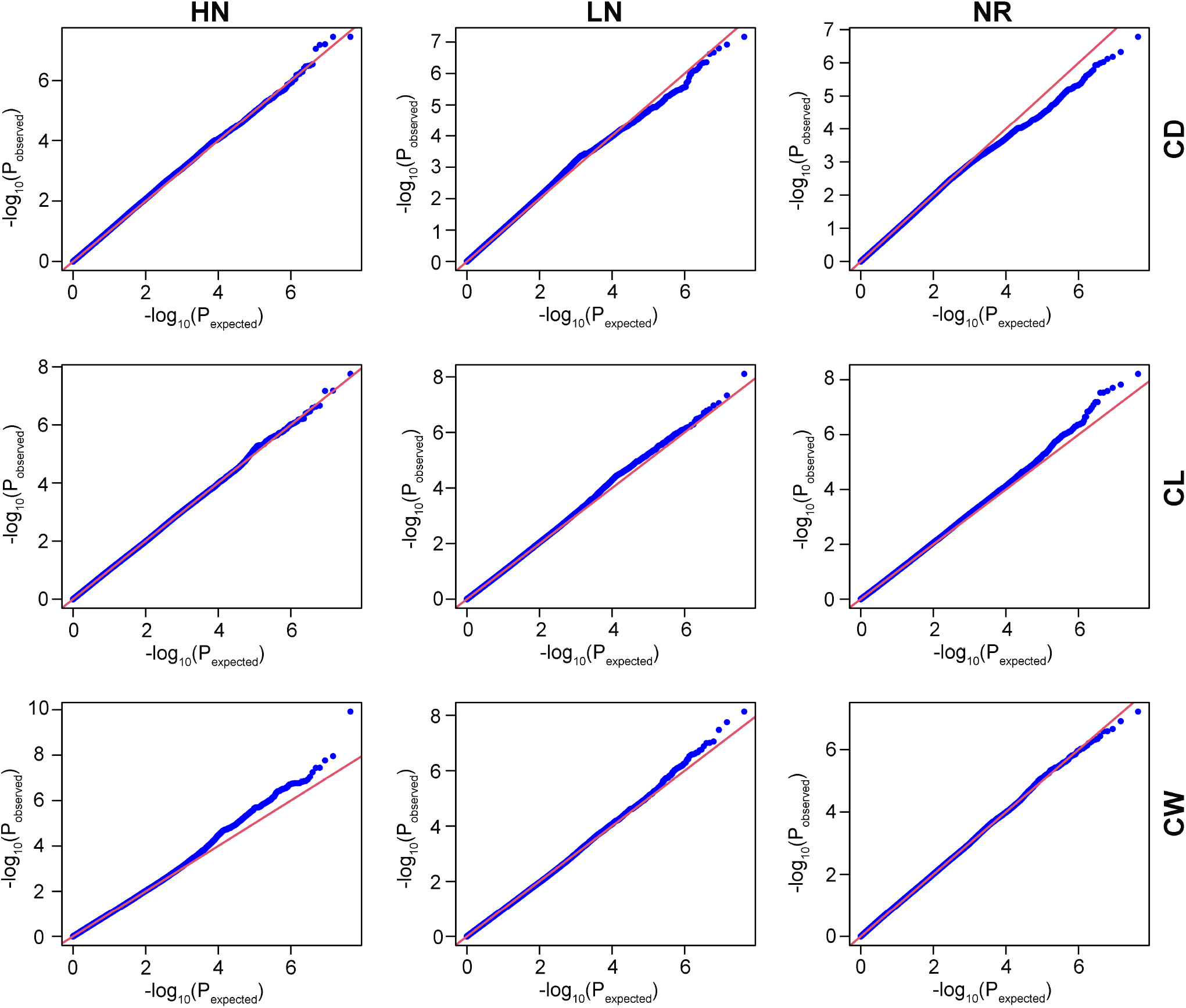
Quantile-quantile (Q-Q) plots for cob-related traits. Red diagonal line indicates the expected values and the blue dots represent the observations.

**Figure S2.**
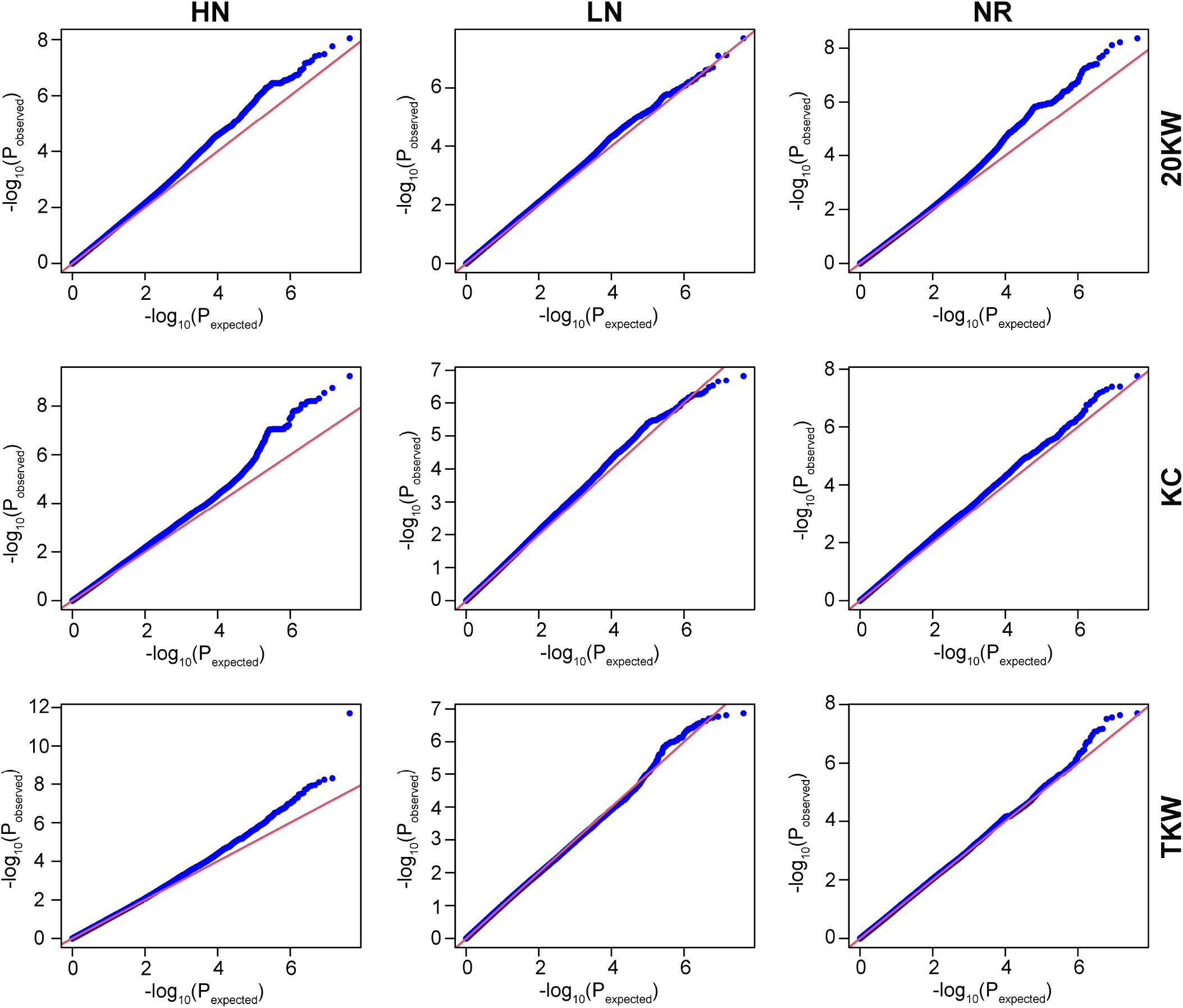
Quantile-quantile (Q-Q) plots for kernel-related traits. Red diagonal line indicates the expected values and the blue dots represent the observations.

**Figure S3.**
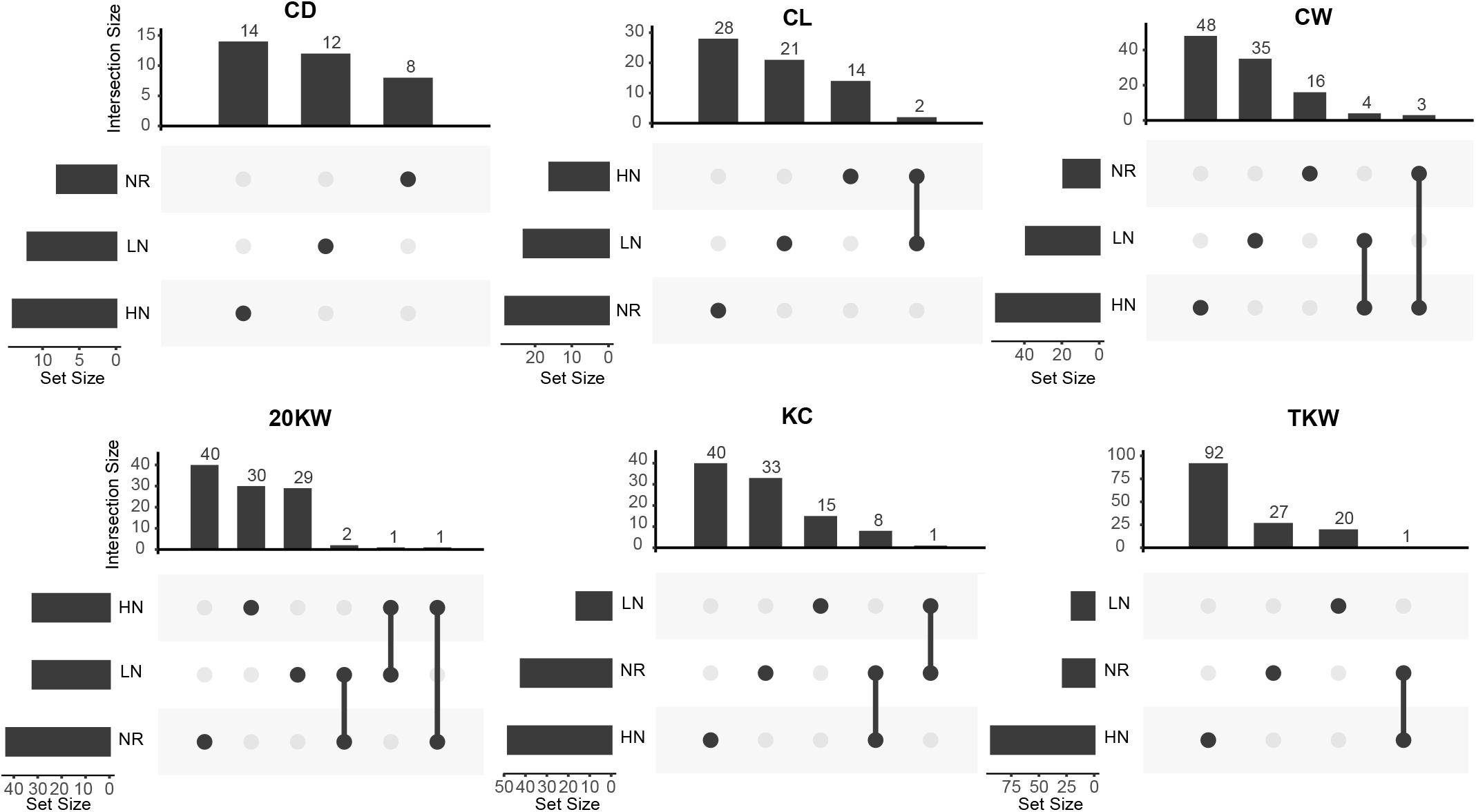
Overlapping results of trait-associated loci (TALs) for each trait under three different N conditions. Numbers on top of the barplots indicate the number of unique (only dots) and shared (dots and lines) TALs.

**Figure S4.**
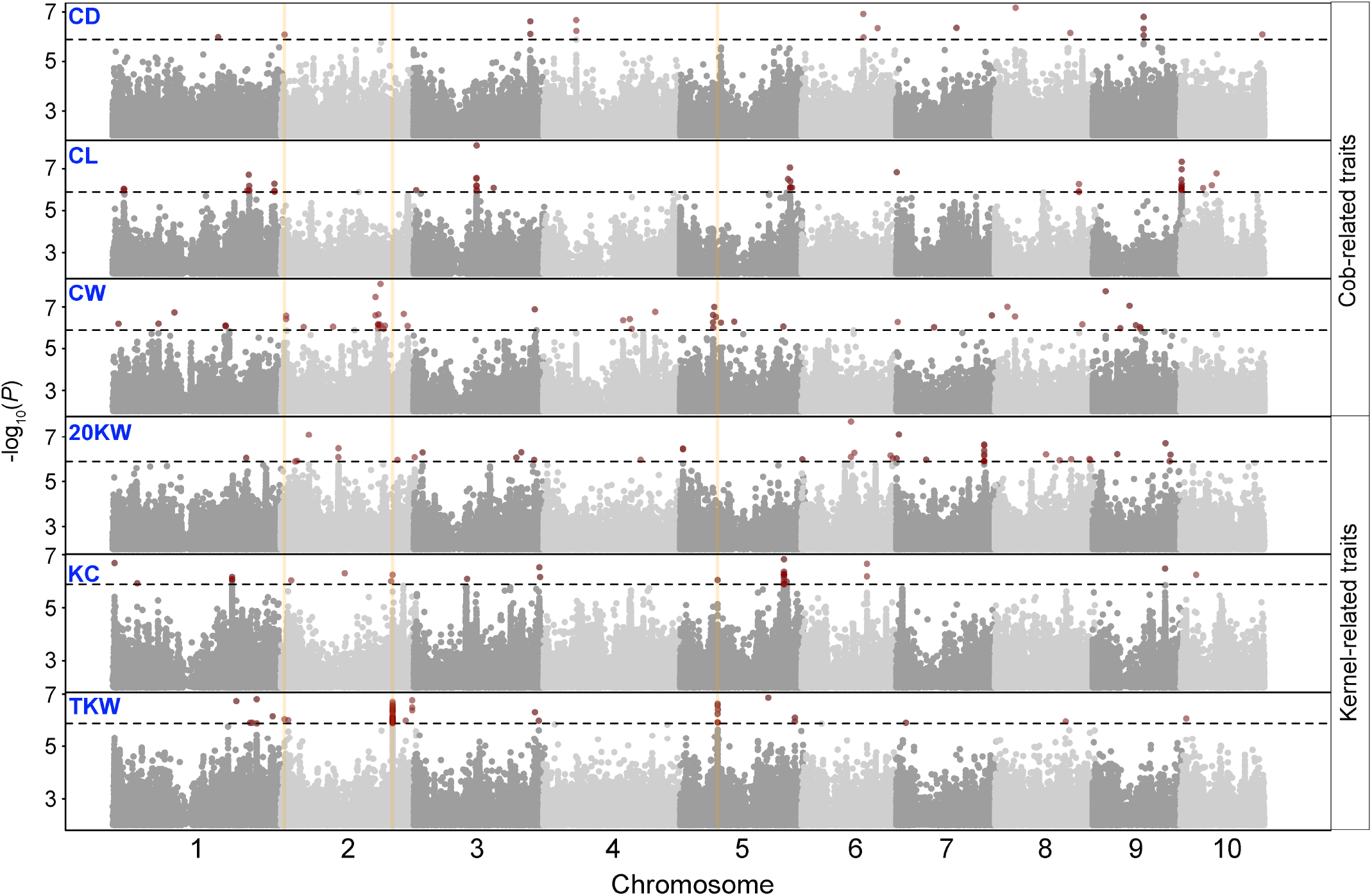
Stacking Manhattan plot of cob- and kernel-related traits under LN conditions. The black horizontal dashed line indicates the GWAS threshold. Each red dot above the threshold represents the SNP significantly associated with a trait. The vertical orange lines indicate the overlapped trait-associated loci (TALs).

**Figure S5.**
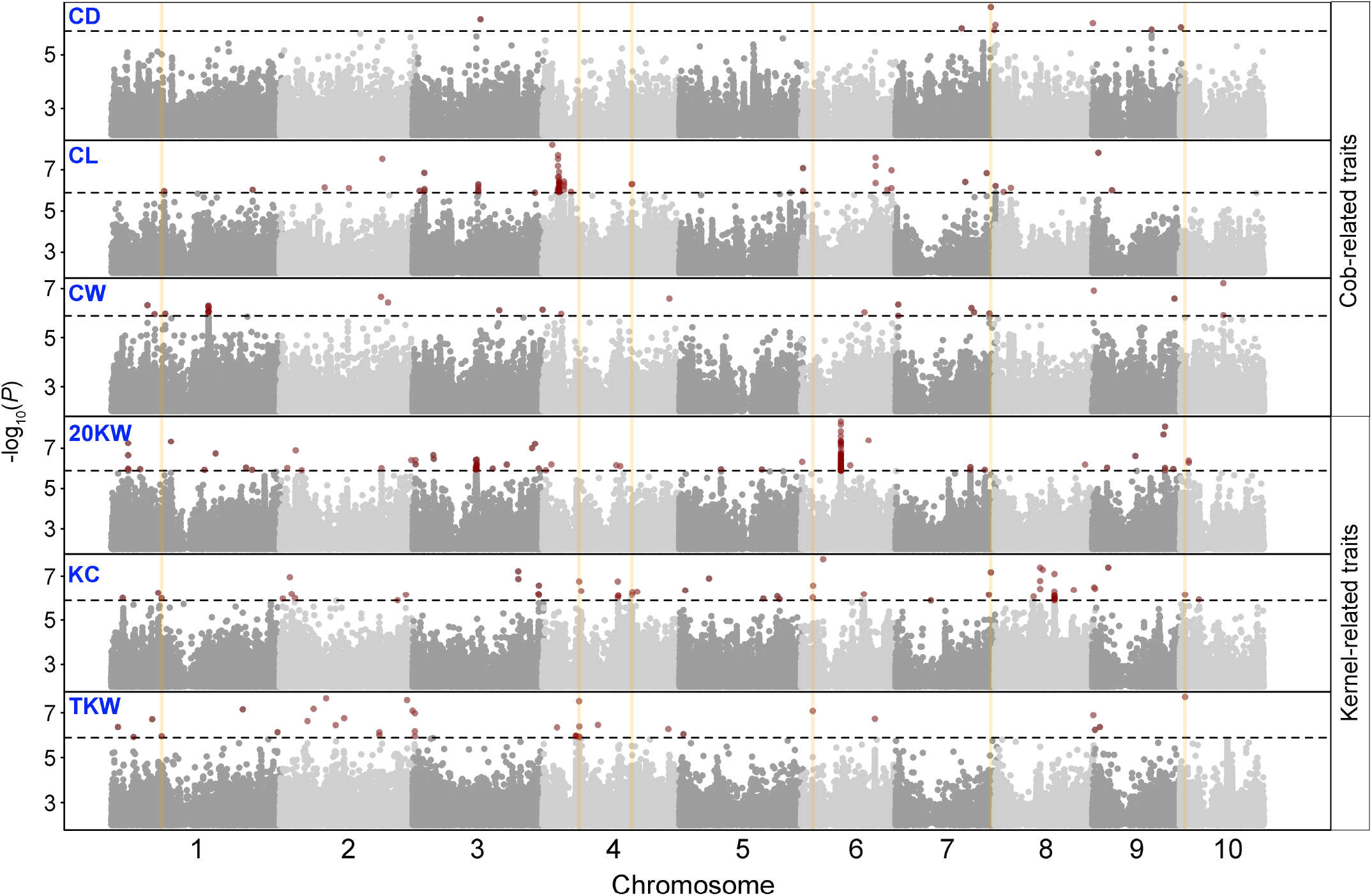
Stacking Manhattan plot for N-responsive (NR) traits. The black horizontal dashed line indicates the GWAS threshold. Each red dot above the threshold represents the SNP significantly associated with a trait. The vertical orange lines indicate the overlapped trait-associated loci (TALs).

## References

1. Pilbeam, D. J. The utilization of nitrogen by plants: a whole plant perspective. Ann. Plant Rev. Online 305–351, DOI: 10.1002/9781444328608.ch13 (2018).

2. Heffer, P. Assessment of fertilizer use by crop at the global level 2010-2010/11 international fertilizer industry association (ifa), paris, france (2013).

3. Ludemann, C., Gruere, A., Heffer, P. & Dobermann, A. Global data on fertilizer use by crop and by country, DOI: 10.5061/dryad.2rbnzs7qh (2022).

4. Gu, B. et al. Abating ammonia is more cost-effective than nitrogen oxides for mitigating pm2. 5 air pollution. Science 374, 758–762 (2021).

5. Agrama, H., Zakaria, A., Said, F. & Tuinstra, M. Identification of quantitative trait loci for nitrogen use efficiency in maize. Mol. Breed. 5, 187–195 (1999).

6. Liu, R. et al. Mining of candidate maize genes for nitrogen use efficiency by integrating gene expression and qtl data. Plant Mol. Biol. Report. 30, 297–308 (2012).

7. Coque, M., Martin, A., Veyrieras, J., Hirel, B. & Gallais, A. Genetic variation for n-remobilization and postsilking n-uptake in a set of maize recombinant inbred lines. 3. qtl detection and coincidences. Theor. Appl. Genet. 117, 729–747, DOI: 10.1007/s00122-008-0815-2 (2008).

8. Li, P. et al. A genetic relationship between nitrogen use efficiency and seedling root traits in maize as revealed by QTL analysis. J. Exp. Bot. 66, 3175–3188, DOI: 10.1093/jxb/erv127 (2015). https://academic.oup.com/jxb/article-pdf/66/11/3175/17136978/erv127.pdf.

9. Morosini, J. S. et al. Association mapping for traits related to nitrogen use efficiency in tropical maize lines under field conditions. Plant Soil 421, 453–463 (2017).

10. He, K. et al. Mining of candidate genes for nitrogen use efficiency in maize based on genome-wide association study. Mol. Breed. 40, 1–17, DOI: 10.1007/s11032-020-01163-3 (2020).

11. Ertiro, B. T. et al. Genetic dissection of nitrogen use efficiency in tropical maize through genome-wide association and genomic prediction. Front. Plant Sci. 11, doi: 10.3389/fpls.2020.00474 (2020).

12. Yang, J. et al. Incomplete dominance of deleterious alleles contributes substantially to trait variation and heterosis in maize. PLoS genetics 13, e1007019 (2017).

13. Charlesworth, D., Charlesworth, B. & Morgan, M. The pattern of neutral molecular variation under the background selection model. Genetics 141, 1619–1632 (1995).

14. Flint-Garcia, S. A. et al. Maize association population: a high-resolution platform for quantitative trait locus dissection. The Plant J. 44, 1054–1064, DOI: https://doi.org/10.1111/j.1365-313X.2005.02591.x (2005). https://onlinelibrary.wiley.com/doi/pdf/10.1111/j.1365-313X.2005.02591.x.

15. Rodene, E. et al. A uav-based high-throughput phenotyping approach to assess time-series nitrogen responses and identify trait-associated genetic components in maize. The Plant Phenome J. 5, e20030. DOI: https://doi.org/10.1002/ppj2.20030 (2022). https://acsess.onlinelibrary.wiley.com/doi/pdf/10.1002/ppj2.20030.

16. Meier, M. A. et al. Association analyses of host genetics, root-colonizing microbes, and plant phenotypes under different nitrogen conditions in maize. eLife 11, e75790, DOI: 10.7554/eLife.75790 (2022).

17. Bates, D., Maechler, M., Bolker, B. & Walker, S. Mixed-effects models using lme4. J Stat Softw 67, 1–48 (2015).

18. Xu, G., Fan, X. & Miller, A. J. Plant nitrogen assimilation and use efficiency. Annu. review plant biology 63, 153–182 (2012).

19. Bukowski, R. et al. Construction of the third-generation Zea mays haplotype map. GigaScience 7, DOI: 10.1093/gigascience/gix134 (2017). Gix134, https://academic.oup.com/gigascience/article-pdf/7/4/gix134/24619986/gix134.pdf.

20. Yu, J. et al. A unified mixed-model method for association mapping that accounts for multiple levels of relatedness. Nat. genetics 38, 203–208, DOI: 10.1038/ng1702 (2006).

21. Parisseaux, B. & Bernardo, R. In silico mapping of quantitative trait loci in maize. Theor. Appl. Genet. 109, 508–514, DOI: 10.1007/s00122-004-1666-0 (2004).

22. Chang, C. C. et al. Second-generation PLINK: rising to the challenge of larger and richer datasets. GigaScience 4, DOI: 10.1186/s13742-015-0047-8 (2015). S13742-015-0047-8, https://academic.oup.com/gigascience/article-pdf/4/1/s13742-015-0047-8/25512027/13742_2015_article_47.pdf.

23. Zhou, X. & Stephens, M. Genome-wide efficient mixed-model analysis for association studies. Nat. genetics 44, 821–824 (2012).

24. Zeng, J. et al. Signatures of negative selection in the genetic architecture of human complex traits. Nat. genetics 50, 746–753, DOI: 10.1038/s41588-018-0101-4 (2018).

25. Yang, Z., Xu, G., Zhang, Q., Obata, T. & Yang, J. Genome-wide mediation analysis: an empirical study to connect phenotype with genotype via intermediate transcriptomic data in maize. Genetics 221, iyac057 (2022).

26. Liu, Y. et al. Genomic basis of geographical adaptation to soil nitrogen in rice. Nature 590, 600–605, DOI: 10.1038/s41586-020-03091-w (2021).

27. Falconer, D. S. & Mackay, T. F. C. Introduction to quantitative genetics (Harlow: Longman, 1996), fourth edn.

28. Yang, Z., Xu, G., Zhang, Q., Obata, T. & Yang, J. Genome-wide mediation analysis: bridging the divide between genotype and phenotype via transcriptomic data in maize. bioRxiv (2021).

29. Li, M.-X., Yeung, J. M., Cherny, S. S. & Sham, P. C. Evaluating the effective numbers of independent tests and significant p-value thresholds in commercial genotyping arrays and public imputation reference datasets. Hum. genetics 131, 747–756 (2012).

30. Ribeiro, P. et al. Identification of quantitative trait loci for grain yield and other traits in tropical maize under high and low soil-nitrogen environments. Crop. Sci. 58, 321–331, DOI: https://doi.org/10.2135/cropsci2017.02.0117 (2018). https://acsess.onlinelibrary.wiley.com/doi/pdf/10.2135/cropsci2017.02.0117.

31. Yang, H. et al. Temporal and spatial variations of soil c, n contents and c:n stoichiometry in the major grain-producing region of the north china plain. PLOS ONE 16, 1–14, DOI: 10.1371/journal.pone.0253160 (2021).

32. Zhu, H., Chen, X. & Zhang, Y. Temporal and spatial variability of nitrogen in rice–wheat rotation in field scale. Environ. earth sciences 68, 585–590, DOI: 10.1007/s12665-012-1762-4 (2013).

